# Pathogen prevalence modulates medication behavior in ant *Formica fusca*

**DOI:** 10.1101/2022.02.09.479685

**Authors:** Jason Rissanen, Heikki Helanterä, Dalial Freitak

**Author notes:** Corresponding author: Jason Rissanen, University of Graz, Institute of Biology, Universitätsplatz 2, 8010 Graz, Austria.

## Abstract

Ants face unique challenges regarding pathogens, as the sociality which has allowed them to form large and complex colonies also raises the potential for transmission of disease within these colonies. To cope with the threat of pathogens, ants have developed a variety of behavioral and physiological strategies. One of these strategies is self-medication, in which animals use biologically active compounds to combat pathogens in a way which would be harmful in the absence of them. *Formica fusca* ants are to date the only species of ants proven to successfully self-medicate against an active infection caused by a fungal pathogen by supplementing their diet with food containing hydrogen peroxide. Here, we build on that research by investigating how the prevalence of disease in colonies of *F. fusca* affects the strength of the self-medication response. We exposed either half of the workers of each colony or all of them to a fungal pathogen and offered them different combinations of diets. We see that workers of *F. fusca* engage in self-medication behavior even if exposed to a low lethal dose of a pathogen, and that the strength of that response is affected by the prevalence of the disease in the colonies. We also saw that the infection status of the individual foragers did not significantly affect their decision to forage on either control food or medicinal food as uninfected workers were also foraging on hydrogen peroxide food, which opens up the possibility of kin medication in partially infected colonies. Our results further affirm the ability of ants to self-medicate against fungal pathogens, shed new light on plasticity of self-medication and raise new questions to be investigated on the role self-medication has in social immunity.

## Introduction

Eusociality has been a key driver in why ants have become some of the most successful insects in the world, inhabiting most terrestrial environments on the planet (1). The high degree of cooperation and the division of labour of ants benefit the colonies in the form of more effective brood care and acquisition and protection of resources, which has allowed ant colonies to grow large, often comprising of hundreds of thousands or even millions of individuals (2). Sociality also comes with costs. Ants live in dense colonies with a low genetic diversity among frequently interacting nestmates, which creates favourable conditions for pathogens to spread (3,4). Ants also forage and interact with a wide variety of organisms in their environment, which makes them vulnerable to exposure to pathogens (5). The combination of high risk of exposure as well as favourable conditions for transmission of pathogens makes ants particularly susceptible for outbreaks of infectious diseases (3).

To cope with the high pathogen pressure they face, a wide variety of both physiological and behavioral strategies have evolved in ants. Ants share an innate immunity with other insects as a first line of defence, but also have unique adaptations in the form of anti-microbial gland exudates (6–8). Ants also rely on behavior to initially avoid pathogens, but if exposed they can mitigate the effects of them through hygienic behaviors such as grooming of both self and nestmates (9). The extreme level of sociality also means that the anti-pathogenic strategies of individual ants in a colony lead to a whole which is larger than the sum of its parts, providing protection against pathogens on a colony level as a form of social immunity (10,11).

As a complementary strategy, ants could potentially use self-medication to stave off the threat of disease. Self-medication is the use of biologically active compounds to fight off disease in a way which would be harmful or costly for uninfected individuals (12). The self-medication response is plastic and can take place either before infection takes place (prophylactic) or after (therapeutic) (13), and can be directed either at self or kin (12,14). To be qualified as true self-medication, the behavior needs to fulfil four criteria: the compound must be deliberately contacted (I), the compound must be harmful for the pathogen (II), the behavior must lead to a higher fitness for infected individuals (III), the use of the compound must be harmful for uninfected individuals (IV) (12).

Evidence of self-medication behavior in ants and other social insects is still scarce (15–17) and to date the clearest evidence for self-medication has been provided in solitary insects (13,18–21). So far, there is only one demonstrated case of therapeutic self-medication in ants (17), where workers infected with a highly lethal dose of a generalist entomopathogenic fungus consumed food laced with hydrogen peroxide (H_2_O_2_), a reactive oxygen species (ROS) which is part of their natural immune responses(22–24), to increase their chances of survival. There is also a case to be made that the collection and processing of resin into the nest material is a form of prophylactic self-medication in ants (8). However, the extent of self-medication responses and how they play out in the colony foraging dynamics is still unknown in ants.

Due to the eusociality of ants, individual self-medication responses are likely to engage colony level processes which regulate the collection and distribution of medicinal compounds in the colony. The division of labour in ant colony means that the foragers, who commonly make up around 10% — 20% of the workers in the colony (1,25–27), provide nutrition for the entire nest, and tune their foraging choices according to the nutritional needs of their nestmates (28). Therefore, if nestmates are battling disease, the nutritional needs of the colony change, and foragers should respond by foraging on medicinal compounds regardless of their individual infection status to distribute medicine to infected nestmates as a form of kin medication (12).

In this study, we tested how different levels of pathogen infection affect the self-medication behavior of *Formica fusca* ants in the form of foraging choices of colonies, and whether they lead to enhanced survival against the pathogen. The use of different levels of pathogen exposure combined with the use of either fixed food diets or a diet choice allowed us to explore questions relating to the plasticity of the self-medication response. We predict that as the prevalence of disease in a colony increases, so does the strength of the self-medication response. By using colonies in which only part of the workers is infected, we could also investigate a potential kin medication component to the self-medication response. We also predict that both infected and healthy workers will forage on medicinal food, with the task of distributing it to the rest of the colony. We used a dose of pathogen which had a low effect on mortality, which is considered to be more naturally relevant (29). Studies using highly lethal doses have also been shown to negatively affect normal immunologic behaviors such as grooming (30).

## Material and methods

We collected 47 wild nests of *Formica fusca* from around the Hanko peninsula in southern Finland (59°54’46.9”N, 23°15’53.8”E). *Formica fusca* is a weakly polygynous species, and some of the nests contained several queens. From the nests we made a total of 133 experimental colonies of one queen and 100 workers each and placed them in separate plastic containers (8 cm x 15 cm x 10 cm). Whenever several experimental colonies were made of a single nest, they were used in the different setups. The walls and lids of each container was lined with a 20% (w/v) mixture of ethanol and talcum powder to keep the ants from escaping (31). A small portion of the lid was cut off to get a clear view of the foraging area of the containers. Each container had a 2 cm deep plaster cast base with a 1 cm deep circular indentation (3 cm Ø) as a nesting site with a ceramic tile (4 cm x 4 cm) on top for shelter.

### Experimental setup

To study how the strength of the self-medication response of a colony changes with increasing disease prevalence, we set up an experiment using three different ratios of infected vs. noninfected workers and exposed all the treatments to three different feeding regimes. This way, we could study how the foraging activity changes on a colony level as well as how the infection status of individual foragers affects their foraging choices.

The 133 colonies were divided into three groups to be used in the three different infection ratios. 44 colonies were used as infection free sham-treated controls (0%), in 45 colonies half of the workers were infected and half sham-treated (50%), in the remaining 44 colonies, all of the workers were infected (100%). Queens were not subjected to any treatment in any of the colonies. To be able to distinguish infected ants from healthy ants in the 50% infected colonies, half of the workers were marked with a red dot on their abdomen using marking colour for marking honeybee queens. The marked individuals in the colonies were subjected to either the infection or control treatment to limit any differences caused by handling. To ensure that the marking would not cause differences in behavior due to handling in relation to the colonies in the other treatments, we marked half the workers in all of the experimental colonies the same way. The marking of ants also functioned as a blinding measure during the observational phase to limit observational bias.

15 colonies of each infection ratio were given two Eppendorf tube caps of the Bhatkar & Whitcomb diet (32) as the control food. 15 colonies were given a food choice of one cap of control food and one cap containing control food containing 4% H_2_O_2_, the same concentration previously used by Bos et al. (17). The remaining 15 colonies were given a diet of two caps containing 4% H_2_O_2_ food (ROS diet). The experimental setup is visualized in figure 1. Fresh food in Eppendorf caps were provided each morning at 11 a.m. Each colony also had an Eppendorf tube with water and cotton to drink from.

**Figure 1.**
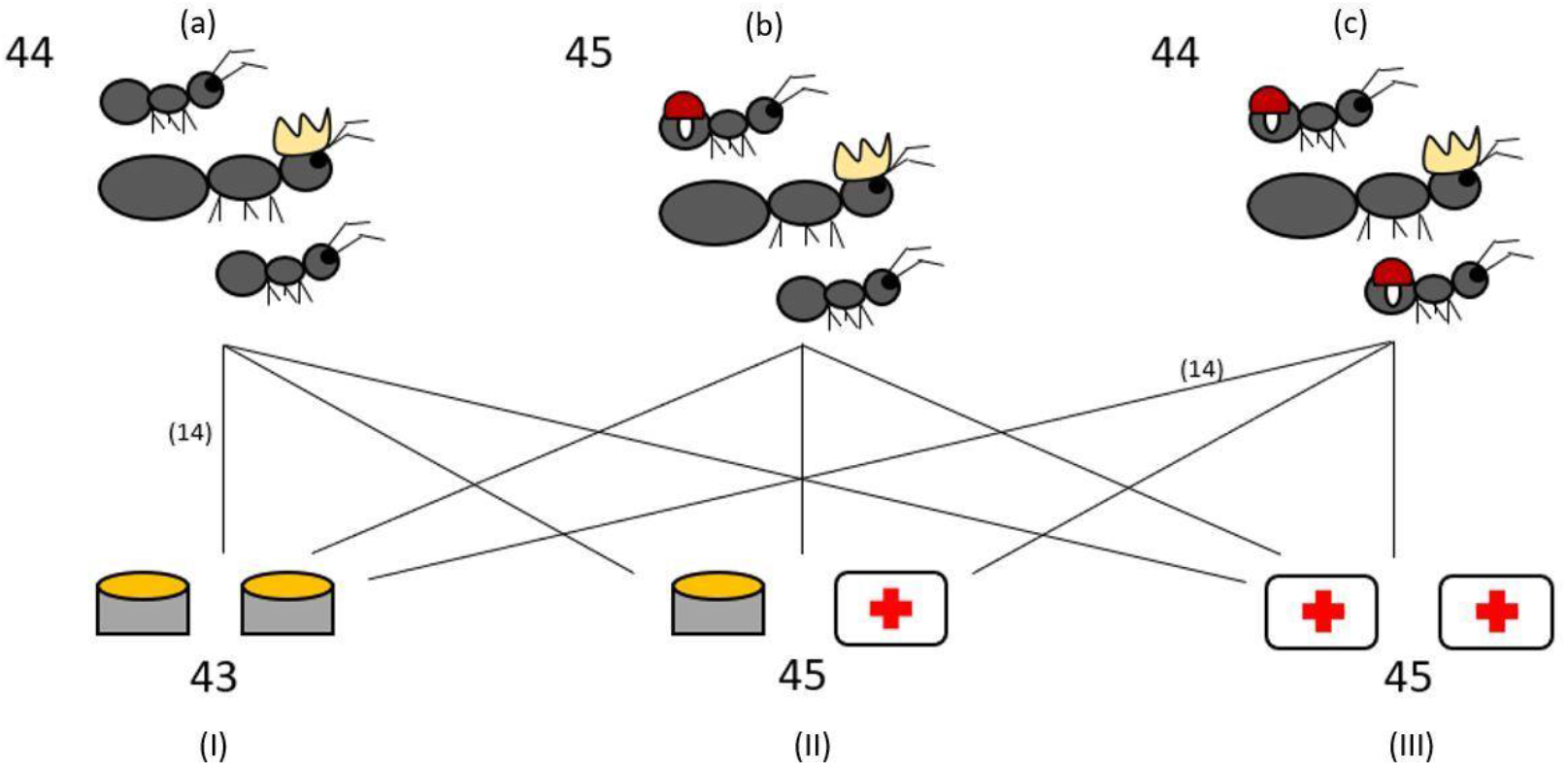
Experimental setup. The 133 experimental colonies were split into three different infection treatments: 0% workers infected (a), 50% workers infected (b), 100% workers infected (c). Colonies from each infection treatment were further assigned to one of three different diets: fixed control food diet (I), food choice diet between control food and 4% H_2_O_2_ food as medicine (II), fixed 4% H_2_O_2_ diet (III). The numbers indicate the total number of experimental colonies used in the different treatments and diets. The lines all represent 15 colonies, with the exception of two groups which have 14 colonies.

The foraging was observed four times a day (10, 14, 18 and 22) for a total duration of six days. At each observation period, the colonies were photographed through the opening in the lid and the foraging was determined by analyzing the photographs. If an ant had contact with the food with any appendage, it was deemed as a forager. The strict criterion for foraging was to prevent other activity to be assessed as foraging. From each photograph, every forager was counted. We also made a note on what food was being foraged and whether the ants were marked or not. Photographs were analyzed blind with respect to the infection and diet treatment.

### Infections

*Beauveria bassiana* was grown on petri dishes with Potato Glucose Agar, incubated in the dark at room temperature. To prevent any effect of local adaptations to *B. bassiana*, we used a Danish strain (KLV 03-90) of fungus which has been previously used with *Formica fusca* (17). Spores of *B. bassiana* were collected from plates with visibly sporulating fungus by pipetting 10mL of 1 x PBS on the plate and then carefully rubbing it with a sterile glass rod to collect spores. The solution of PBS containing spores was centrifuged in 3000 rpm for 3 minutes in 4°C. The supernatant was discarded and the spores were then suspended in 10mL of Milli-Q H_2_O. The spore concentration was determined using a haemacytometer.

To infect the ants, we submerged them in a solution containing 1 x 10^7^ spores/mL of *B. bassiana* for five seconds. The control ants were all submerged in MilliQ-H_2_O for five seconds. The colonies were given a day to settle in their nests, with both water and control food being provided, before we started to observe foraging behavior. Dead ants were not removed during the observation period to keep the pathogen pressure on the colonies high.

### Statistical analysis

All statistical analysis was done using the R software (version 4.1.2, R Core Team 2020).

The foraging data was analyzed using generalized linear mix models (GLMM) using the glmmTMB function from the *glmmTMB* package (33). All the models were fitted with a poisson distribution and were tested for deviations in dispersion using the *DHARMa* package for diagnostics for hierarchical regression models (34). Original nest and time nested within the experimental colony ID were used as random effects in all the models to account for nest-caused differences as well as pseudoreplication. Pairwise comparisons were conducted using the emmeans function from the *emmeans* package (35) using a Tukey’s p-value adjustment when performing multiple comparisons.

The model for analyzing overall foraging activity used the number of ants foraging for food during the observational periods as the response variable. The infection treatment (0%/50%/100%) and diet (fixed control/food choice/fixed ROS) were used as fixed factors, as well as the interaction between them.

Foraging within the food choice diet was analyzed with a model using the total number of ants foraging on the different foods during the observational periods as the response variable, with the infection treatment and food type (control food/ROS food) and their interaction as fixed factors.

Whether the infection status of the individuals affects the choice on foraging on the different food types in the food choice diet was analyzed with a model using the total amount of ants foraging on the different foods within the 50% infection treatment as the response variable with the infection status of the individuals (infected/uninfected) and food type and their interaction as fixed factors.

The survival data was analyzed using a cox proportional hazard model from the *coxme* package (36). The numbers of individual ants who had died when the observation was terminated was used as the response variable, and the infection treatment and diet, as well as their interaction, were used as fixed factors. Original nest was used as a random effect to account for within nest differences. The experimental colony was also used as a random effect to account for pseudoreplication. Pairwise comparison between the groups was done using the emmeans function with a Tukey’s p-value adjustment for multiple group comparisons.

Experimental colony 14 experienced abnormally high mortality and was therefore omitted from the analysis. The queen died during the observational period in experimental colonies 70 and 75 and were omitted from the analysis due to possible alterations in the natural behavior of the colonies caused by death of the queen. The omission of these colonies did not affect the results in a significant way.

## Results

### Foraging

Access to the different diets had a significant effect on foraging activity in the colonies (foraging ^~^ diet, X^2^_2_ = 20.868, p < 0.001), but the different infection treatments did not (foraging ^~^ infection treatment, X^2^_2_ = 3.963, p = 0.138). The interaction between infection treatment and diet was not significantly affecting the overall foraging activity of the colonies (infection treatment × diet interaction on foraging, X^2^_12_ = 0.386, p = 0.984). Ants were foraging significantly less on a fixed ROS diet compared to both the control diet (t = 3.801, p < 0.001) and the food choice diet (t = 4.157, p < 0.001), but there was no difference in foraging activity when colonies had access to the control and food choice diets (t = −0.282, p = 0.957) (Figure 2).

**Figure 2.**
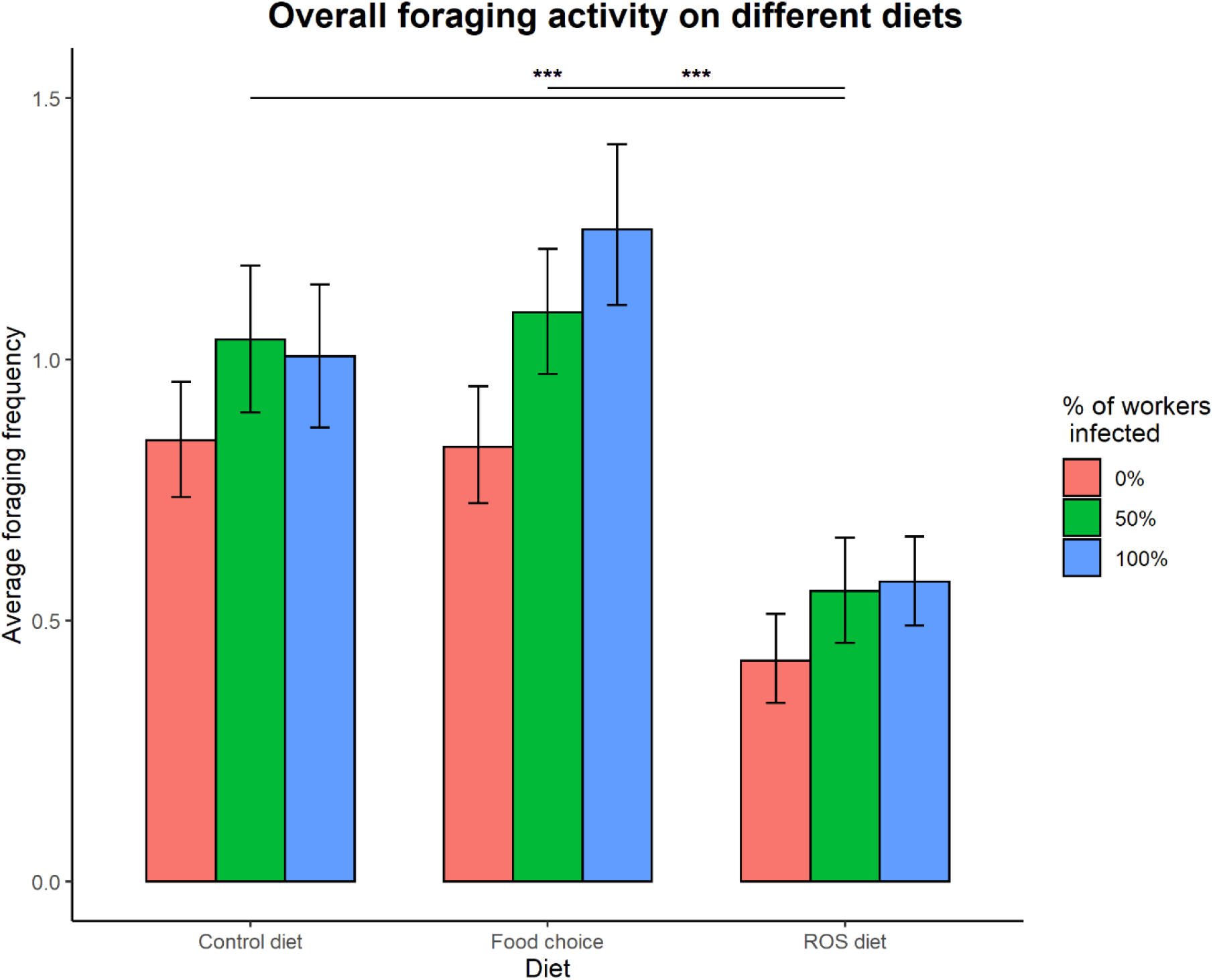
Overall foraging activity on different diets. Access to the different diets affected foraging activity in the colonies (X^2^_2_ = 19.189, p < 0.001), as foraging on a fixed ROS diet was significantly lower compared to both the control diet (t = 3.781, p < 0.001) and the food choice diet (t = 4.129, p < 0.001). The difference between foraging on the control diet and the food choice diet was not significant (t = −0.277, p = 0.959). Infection treatment did not affect foraging in a significant way (X^2^_2_ = 3.865, p = 0.149). The error bars represent the 95% confidence intervals.

Within the food choice diet, the effect of infection treatment on foraging activity depended on the type of food (control food vs. ROS food) (infection treatment × food type interaction on foraging, X^2^_2_ = 8.025, p = 0.018). Pairwise comparisons revealed that foraging on control food within the food choice diet was not affected by infection treatment in colonies (0%:50%, t = −1.095, p = 0.518; 0%:100, t = −0.995, p = 0.580; 50%:100%, t = 0.113, p = 0.993) (Figure 3A). However, foraging activity on ROS food was affected by the disease prevalence when presented with a food choice diet. 100% infected colonies were foraging significantly more on ROS food compared to 0% infected colonies (t = −2.873, p = 0.012), and the differences in foraging between the 0% and 50% or the 50% and 100% colonies were not significant (0%:50%, t = −2.873, p = 0.251; 50%:100%, t = −1.311, p = 0.389) (Figure 3B). Foraging on control food compared to ROS food was much more common regardless of the infection treatment of the colonies (0%, t = 11.272, p < 0.001; 50%, t = 12.352, p < 0.001; 100%, t = 13.045, p < 0.001).

**Figure 3.**
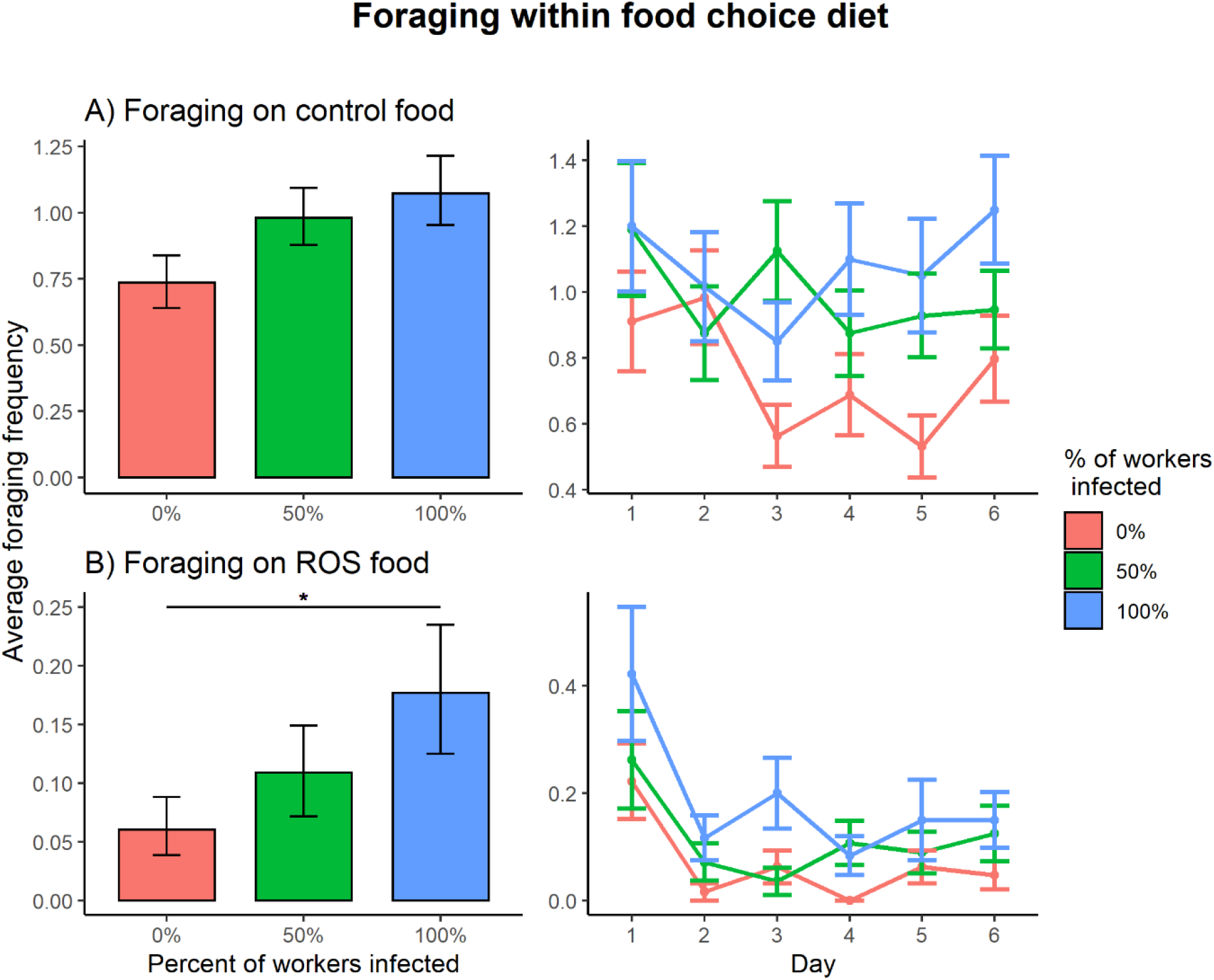
Foraging frequencies between the infection ratios 0% (red), 50% (green), 100% (blue) within the food choice setup. (A) There were no significant differences in foraging activity on control food between any of the infection treatments. (B) When foraging on ROS food, the 100% infected colonies were foraging significantly more compared to the 0% infected colonies (t = −2.873, p = 0.012), but the differences between 0%:50% and 50%:100% infected colonies were non-significant. The error bars represent the 95% confidence interval.

Infection status did not significantly affect the overall foraging activity (foraging ^~^ infection status, X^2^_1_ = 1.527, p = 0.217). Foraging activity on the different types of food (control food vs. ROS food) in the 50% colonies with access to a food choice was not dependent on the infection status (uninfected vs. infected) of the individuals (infection status × food type interaction on foraging, X^2^_1_ = 2.694, p = 0.101) (Figure 4).

**Figure 4.**
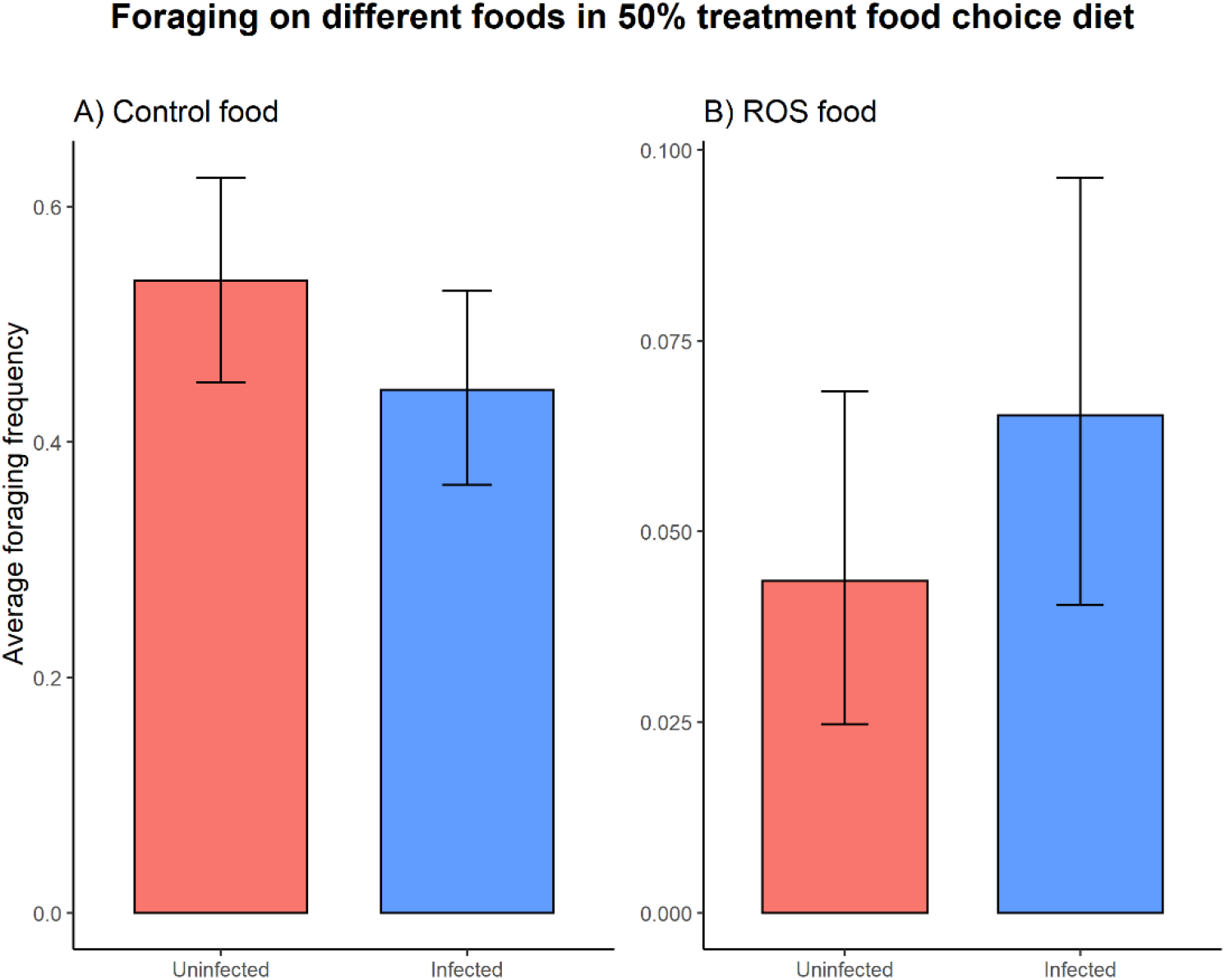
The infection status of the individual did not cause significant differences in foraging activity on either control food (t = 1.685, p = 0.092) (A) or ROS food (t = −1.175, p = 0.240) (B) when presented with a food choice diet. The error bars represent the 95% confidence interval.

### Survival analysis

Both the diet and the infection treatment in the colonies significantly affected the worker mortality in the colonies (diet, X^2^_2_ = 12.802, p = 0.002; infection treatment, X^2^_2_ = 9.148, p = 0.010), but the interaction of them did not affect mortality significantly (diet × infection treatment interaction on survival, X^2^_4_ = 3.006, p = 0.557).

Infecting of all the workers in the colony with *B. bassiana* significantly affected the mortality compared to healthy colonies when no medicinal food was present (0%:100%, z = −2.638, p = 0.023). Infecting half of the workers in the colonies had no significant effect on survival compared to either healthy colonies (0%:50%, z = −1.058, p = 0.540) or colonies where all the workers were exposed (50%:100%, z = −1.591, p = 0.249) (Figure 5A).

**Figure 5.**
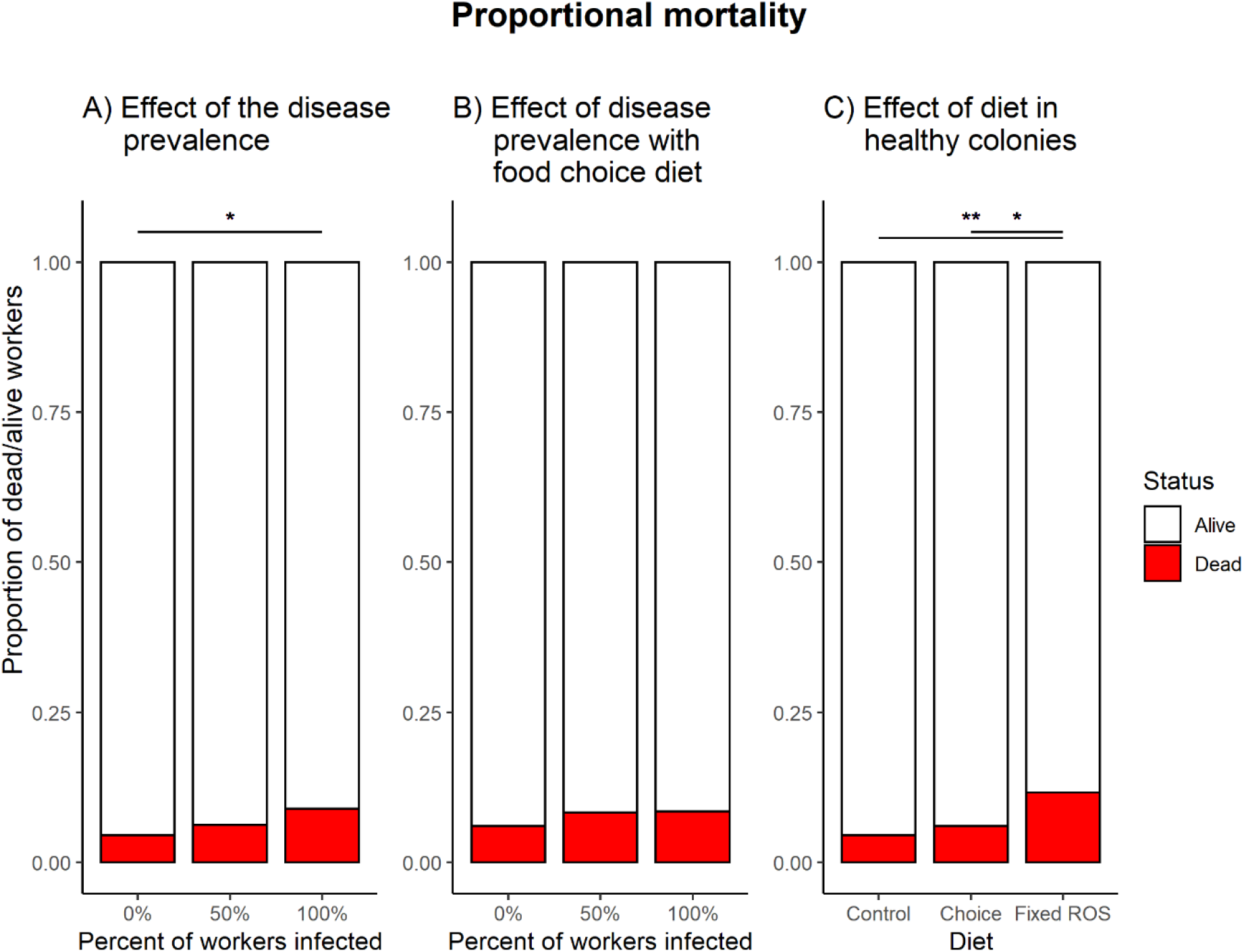
Proportional mortality in the different treatments. (A) Effect of the infection ratio on mortality in colonies fed with the control diet. The dose of pathogen used for the experiment had an overall low effect on mortality, but enough to cause a significant effect on mortality when all of the workers in the colony were infected with *B. bassiana* (z = −2.638, p = 0.023). (B) The effect of infection ratio in colonies with a food choice diet. When access to both foods, the 100% infected colonies no longer show a significantly higher mortality compared to control colonies (z = −1.456, p = 0.312). (C) The effect of the different diet treatments on mortality in healthy colonies. Colonies feeding on a fixed ROS diet had significantly higher mortality compared to colonies feeding on the control diet (z = −3.205, p = 0.004) or the food choice diet (z = −2.403, p = 0.043).

If the ants had access to the food choice diet with both control food and medicinal food, the ratio of workers infected with *B. bassiana* did not affect survival (0%:50%, z = −0.889, p = 0.648; 0%:100%, z = −1.456, p = 0.312; 50%:100%, z = −0.524, p = 0.860) (Figure 5B).

Healthy colonies feeding on a fixed ROS diet have significantly higher mortality compared to colonies who have a control diet (z = 3.205, p = 0.004) or a food choice diet (z = −2.403, p = 0.043). No difference in mortality is observed between healthy colonies with the control diet or having a choice between the two types of food (z = −0.902, p = 0.639) (Figure 5C).

## Discussion

*Formica fusca* ants have previously been shown to self-medicate by incorporating food containing ROS in the form of H_2_O_2_ into their diet in response to a highly lethal dose of *B. bassiana* (17). Here we show that a similar response is also induced with a much less lethal dose, and that the strength of self-medication response is modulated by the ratio of infected ants in the colony, as the colonies were choosing to forage increasingly on ROS food as disease prevalence increased in the colonies, as we predicted.

Adjusting foraging on ROS food according to pathogen prevalence implies plastic and deliberate use of the compound, which is in accordance with the first criterion of self-medication. Our experiment also clearly shows that another criterion is met: ROS food is harmful for healthy individuals. Bos et al. (17) have previously confirmed a further criterion, that H_2_O_2_ lowers the fitness of *B. bassiana*. The final criterion we cannot clearly prove in our experiment – *i.e*. the benefit of the dietary change as a response to the infection. We do see however, that when having access to a balanced diet of both medicinal and normal food, the level of infection does not affect the survival of ants in a significant way compared to healthy colonies, so there is some evidence for a benefit to changing the diet.

Two factors may have masked any benefits in survival caused by changes in foraging behavior: the low effect of the pathogen dose on ant mortality and the relatively short period of time observing the colonies. Observing mortality after seven days gives us only a view on the effects of the primary infection, but the disease could still linger in the colonies due to community transfer of the disease, as dead ants were not removed from the colonies. Therefore, it is reasonable to expect, that the colony would continue to forage on medicinal food which due to its deleterious effects on survival could render short-term benefits low or unclear, but in a longer timeframe benefit the colonies in either worker survival or other measures of fitness. The issue of determining costs and benefits on more than just the short-term survival of individual ants has been raised before (37) and remains an important part in future studies on self-medication behavior.

An interesting result of the experiment was that also infection free colonies with a food choice diet were observed foraging on ROS food, even if this was rarer than in infected colonies. The reason for this behavior is speculative, but it could be possible that the ants are displaying a fixed prophylaxis behavior. de Roode & Lefèvre (38) argued, that if insects would face constant pathogen pressure, evolution would favour prophylaxis to become a fixed behavior. Although this experiment was not enough to provide strong enough evidence for it, Bos et al. (17) reported a similar observation of healthy colonies incorporating ROS food into their diet even if they were free of active infection. As ants are under constant threat of disease due to their behavior and the prevalence of pathogens in their nest and surroundings (3,5,39,40), a fixed prophylaxis would be a feasible strategy for ants to stave off disease, and an interesting study into the extent of the self-medication behavior in ants.

Our results also suggest that obtaining ROS even when only part of a colony is exposed to pathogen infection might not only be an individual response, but instead a response on the colony level. Foraging on H_2_O_2_ supplemented food in the 50% infected colonies was done by both uninfected and infected ants without showing any significant difference due to the health status of the foragers. As the foraging frequency was quite low overall, it is likely that all individual ants weren’t individually foraging on the foods, but instead the foragers of the colony were distributing the food according to the needs of the colony. Buffin et al. (41) have previously shown how food is effectively distributed in colonies of *F. fusca* in a matter of a few hours. If the foragers are distributing medicinal food to infected nestmates as a form of therapeutics as well as uninfected nestmates as prophylaxis, then there could also be a kin medication aspect to the self-medication behavior of ant colonies

In conclusion, this experiment adds to the evidence of the ability of ants to self-medicate against disease. In our experiment, we found that the strength of the self-medication response is plastic and increases when a larger part of the colony is infected with a fungal pathogen. This experiment also takes into consideration the criticism of past experiments, where highly lethal doses of pathogens have been used, to provide a more natural level of exposure of pathogens. The fact that ants alter their foraging behavior in response to not only to prevalence of pathogens in their colonies, but also the quality of the medicinal food available and possibly the severity of the disease (17), means that ants have a highly complex understanding of their surroundings and changes in it caused by pathogens leads to intricate decisions on a colony level to respond to them.

The response of uninfected individuals in a colony to also forage on ROS food implies that there is simultaneous prophylactic as well as therapeutic self-medication within colonies. Thus, the self-medication response is active on both the individual as well as the colony level and opens up the possibility of active kin medication in colonies suffering from pathogen infection. This sort of kin medication also draws parallels with mass drug administration strategies (MDA) used by humans against diseases such as malaria (42,43). In MDA, the entire population is treated against a prevalent disease regardless of whether they are actively infected or not, meaning that simultaneous therapeutic and prophylactic medication takes place, and it has been shown to be an efficient strategy to contain and eliminate the disease (42). Therefore, kin medication driven by foragers could also be a strategy also used by ants to combat pathogens as a part of their social immunity, which opens up an interesting avenue of self-medication research in ants as well as other social insects.

## Author contributions

JR developed the idea for the experiment. JR, HH and DF designed the experiment and carried it out. JR analyzed the data and provided the first draft of the manuscript. All authors worked on the revisions of the manuscript and approved the final version.

## Acknowledgments

We thank Konsta Kesti and Laura Kares for their help with the field work and experiments.

